# Prediction of adverse drug reactions associated with drug-drug interactions using hierarchical classification

**DOI:** 10.1101/2021.02.10.430512

**Authors:** Catherine Kim, Nicholas Tatonetti

## Abstract

Adverse drug reactions (ADRs) associated with drug-drug interactions (DDIs) represent a significant threat to public health. Unfortunately, most conventional methods for prediction of DDI-associated ADRs suffer from limited applicability and/or provide no mechanistic insight into DDIs. In this study, a hierarchical machine learning model was created to predict DDI-associated ADRs and pharmacological insight thereof for any drug pair. Briefly, the model takes drugs’ chemical structures as inputs to predict their target, enzyme, and transporter (TET) profiles, which are subsequently utilized to assess occurrences of ADRs, with an overall accuracy of ~91%. The robustness of the model for ADR classification was validated with DDIs involving three widely prescribed drugs. The model was then applied for interstitial lung disease (ILD) associated with DDIs involving atorvastatin, identifying the involvement of multiple targets, enzymes, and transporters in ILD. The model presented here is anticipated to serve as a versatile tool for enhancing drug safety.

## INTRODUCTION

Adverse drug reactions (ADRs) represent a significant threat to public health worldwide, accounting for considerable morbidity and mortality with estimated costs of ~$500 billion annually [1, 2]. As ADRs continue to present a growing concern in modern health care systems, their identification and prevention are quintessential for improved drug safety and patient care. While drugs are subjected to preclinical *in vitro* safety profiling and clinical drug safety trials to assess drug safety, many ADRs occur in small subsets of the human population, making ADRs not readily detectable in advance [3]. Moreover, ADRs are more difficult to analyze when multiple, rather than single, drugs are administered, which has become common amongst a growing elderly population [4]. Drug-drug interactions (DDIs) between co-administered drugs appear in various forms of ADRs by different mechanisms, adding additional complexity [3].

To better address DDI-associated ADRs, an understanding of their pharmacological mechanisms is strongly required. DDIs can occur when drugs compete for the same target [5]. DDIs also involve drug metabolizing enzymes (e.g. cytochrome P450 (CYP) enzymes) and influx and efflux drug transporters — all of which determine the adsorption, distribution, metabolism, and excretion (ADME) of drugs [6]. Thus, interference with target binding, enzyme-mediated metabolization, and/or uptake and excretion of drugs may cause DDIs [5, 7–9]. Moreover, the comprehensive evaluation of entire TET profiles — many of which are dependent on the chemical structures of drugs — and their interplay between the drugs is critical for an enhanced understanding of DDI-associated ADRs [10].

With the current inability to reliably assess DDIs in preclinical testing and clinical trials and the complex nature of DDI-associated ADRs, a data-driven computational approach is well-suited for predicting such ADRs. This approach may benefit from extensive ADR databases, such as the FDA Adverse Event Reporting System, where data representative of a large population are collected from patients, clinicians, and pharmaceutical companies [11]. While various machine learning models have been previously developed for predicting DDI-associated ADRs with considerable accuracy, they suffer from major limitations. Most currently available models are based on drug similarity, providing accurate prediction only when the drug in question is similar to existing drugs with known TET profiles and/or ADR information [12–15]. This requirement makes these models not readily applicable when such information is unavailable, for example, when a drug is still under development. Moreover, conventional models provide no pharmacological insight into DDI-associated ADRs. The availability of such *a priori* mechanistic understandings can lay out theoretical foundations on which a novel, effective pharmacological strategy can be developed. Overall, a novel computational approach to evaluate associations between DDIs and ADRs and to determine their molecular basis is urgently needed for better drug design and enhanced drug safety.

This study reports the development of a hierarchical machine learning model to predict risks of various DDI-associated ADRs and their underlying pharmacological mechanisms. This model consists of two layers of classifiers for the prediction of TET profiles and occurrences of ADRs from chemical structures of a drug pair, requiring no drug similarity. The model was tested for its robustness with three case studies and then employed to elucidate the origin of an ADR of a rare disease, interstitial lung disease (ILD), associated with DDIs.

## METHODS

All computations were performed with Python 3.8.3 on Jupyter Notebook 6.0.1 Anaconda Navigator 1.9.12, unless otherwise noted.

### Statistical Analyses of ADRs

Proportional reporting ratios (PRRs) were calculated for each drug pair and corresponding adverse drug reactions (ADRs) from the TWOSIDES v0.1 database [3] using the equation 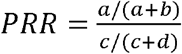 as described previously [3, 16]: where *a* = the number of patients who were administered the drug pair and were reported for the ADR, *b* = the number of patients who were administered the drug pair and were not reported for the ADR, *c* = the number of patients were not administered the drug pair and were reported for the ADR, and *d* = the number of patients who were not administered the drug pair and were not reported for the ADR. The TWOSIDES v0.1 database was created by application of propensity score matching to the FDA Adverse Event Reporting System [11] in order to account for covariates in the dataset and eliminate potential bias [3] and used directly for the PRR calculations in this study. The numbers of unique drug pairs and ADRs used in this study were 211,990 and 12,726, respectively.

For drug pairs containing one of three widely prescribed drugs — levothyroxine, omeprazole, and atorvastatin — all of their reported ADRs were extracted from the TWOSIDES v0.1 database using the Python pandas 1.1.2 library [17].

### Determination of Chemical Fingerprints of Drugs

For all the drugs listed in the DrugBank 5.1.7 database [18], their chemical structures in the format of the simplified molecular-input line-entry system (SMILES) were obtained directly from the database or PubChem v1.6.3.b [18, 19]. The SMILES were stored in a 2D representation with the Python RDKit 2020.03.1 library and used to produce a chemical fingerprint for each drug by calculating its Molecular Access System (MACCS) keys [20]. Binary string representations of the MACCS keys were stored in a Python pandas 1.1.2 dataframe [17].

### Construction of Target, Enzyme, and Transporter Profiles of Drugs

Annotations about 4,263 unique targets, 316 unique enzymes, and 286 unique transporters were collected from DrugBank 5.1.7 [18] to create TET vectors of drugs using the Python NumPy 1.19.4 library [17]. Each of all unique TETs was assigned a position in a TET vector. For each drug, a TET vector was created with the Python NumPy 1.19.4 library to represent its pharmacological profile. Briefly, in each position of a drug’s TET vector, the value of “1” was assigned if any action of the drug (e.g., as a ligand, substrate, inhibitor, activator, agonist, antagonist) on each target, enzyme, and transporter was noted in DrugBank 5.1.7, whereas the value of “0” otherwise.

### Development of RFCs for Prediction of Target, Enzyme, and Transporter Profiles of Drugs

Random forest classifiers (RFCs) were constructed for prediction of targets, enzymes, and transporters from the chemical structure of a drug using the Python sci-kit learn 0.23.2 library [17]. These models formed the first layer of the hierarchical model. The dataset of the drugs’ MACCS keys and TET vectors were split into training (75% of dataset) and testing (25% of dataset) sets. RFCs were trained and tested to predict TET profiles from MACCS key representations (i.e., chemical fingerprints) of the drugs. During training and testing of the RFCs, model accuracies were measured and averaged.

### Development of a Model for Prediction of DDI-associated ADRs from TET Profiles of Drugs

TET vectors of a drug pair were combined to form its TET matrix, which was then matched to the drug pair’s PRRs for various ADRs reported in the TWOSIDES v0.1 database. From the TWOSIDES v0.1 database, the calculated PRRs for different ADRs of each drug pair were categorically encoded with the value of “0” when 0 ≤ PRRs < 1, “1” when PRRs = 1, and “2” when PRRs > 1. The processed PRR dataset with the matched TET matrices for the drug pairs were split into training (75% of dataset) and testing (25% of dataset) sets. The machine learning algorithms, Random Forest Classifiers (RFC) [21], Logistic Regression (LR) [22], and Support Vector Machines (SVM) [23], were constructed as classifiers for ADR prediction using the Python sci-kit learn 0.23.2 library. The models were fit with default tuning parameters in the Python sci-kit learn 0.23.0 library. Model accuracies were measured using a 10-fold cross-validation, as described elsewhere [24]. The SVM model was chosen as a second layer of the hierarchical model.

### Pathway Analysis of ADRs

The key genes/proteins involved in ILD associated with DDIs involving atorvastatin were determined from the pathway database Reactome [25]. Various repositories on gene/protein interactions and pathways, such as BioGRID [26], Proteomics DB [27], STRING [28], and CORUM [29], were applied to identify interactions between drug targets and genes/proteins involved in ADRs. The NCBI Gene database [30] was used to determine tissue-specific gene expression levels.

## RESULTS AND DISCUSSION

### Model Overview

A hierarchical model to predict ADRs from the chemical structures of a drug pair was developed. In this model, the input variables were the chemical structures of drugs (**Fig. 1A**) and the output variables were the PRRs for various ADRs (**Fig. 1B**). A value of PRR > 1 indicates a high risk of an ADR for a given drug pair, whereas a PRR<1 suggests that a given ADR is less commonly reported for a drug pair, relative to other drug pairs [3, 16]. The PRR of 1 indicates a statistically neutral association between a drug pair and an ADR. Instead of attempting to correlate chemical structures of drugs directly with PRRs for ADRs, an intermediate tier of a pharmacological profile, namely a target, enzyme, and transporter (TET) profile, of drugs was introduced to connect chemical fingerprints of drugs with various ADRs (**Fig. 1**).

**Fig. 1.**
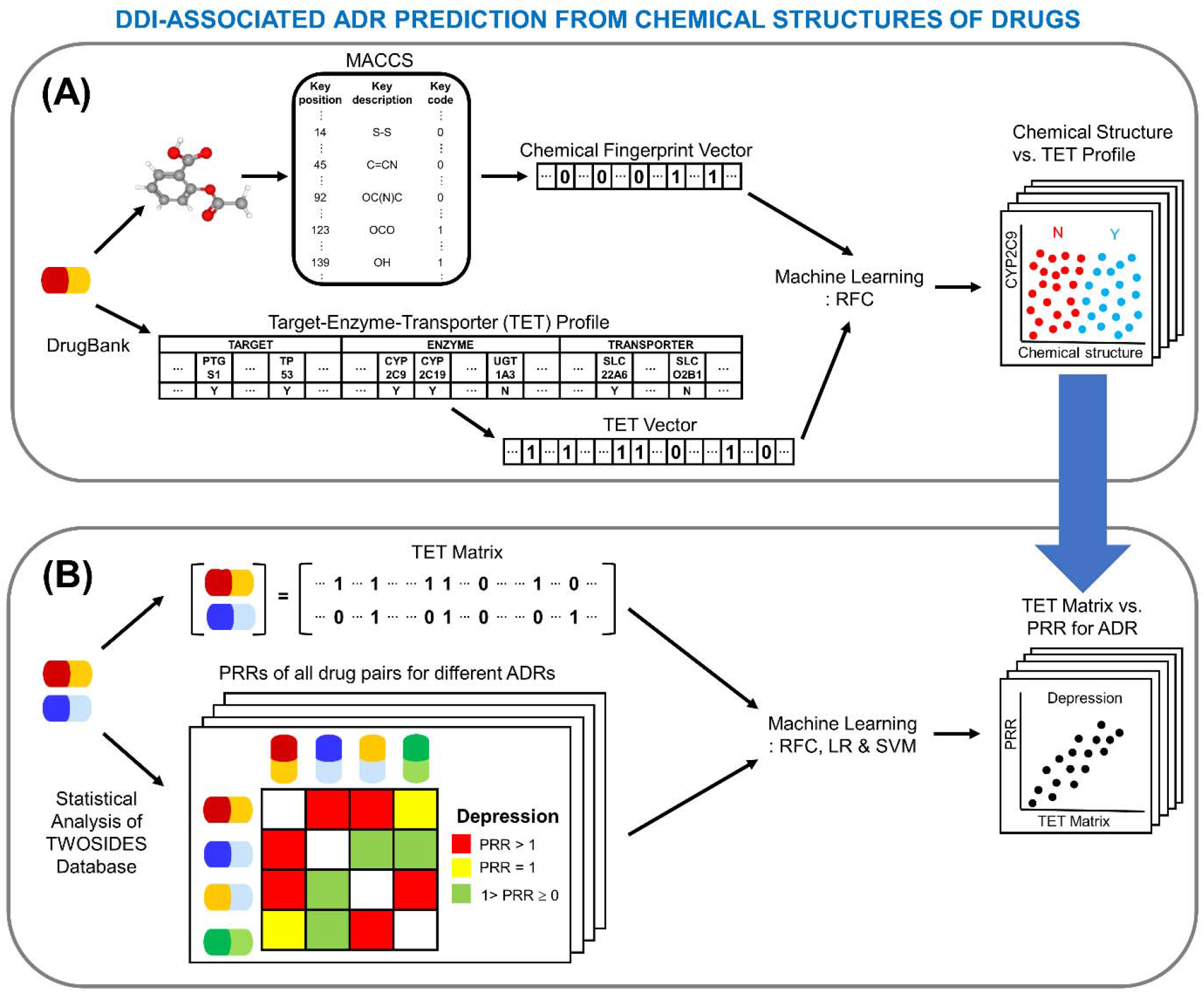
Hierarchical classification model overview for the prediction of DDI-associated ADRs from drugs’ chemical structures through predictions of (A) TET profiles from chemical fingerprints and (B) ADRs from TET matrices of drug pairs. (A) Drugs’ chemical structures were represented with MACCS keys and used as features to predict TET profiles of drugs using a Random Forest Classifier (RFC). (B) TET profiles of a drug pair were combined into a TET matrix, which was then used as a feature to predict encoded PRRs for all ADRs in RFC, Logistic Regression (LR), and Support Vector Machine (SVM) models.

The TET profiles depend on the drugs’ chemical structures [24, 31]. On the other end, the TET profiles determine the drugs’ ADME processes and their ultimate pharmacological actions through on- and off-targeting, all of which play a dominant role in ADRs [5–9]. Thus, the intermediate tier of the TET profiles serves as an essential component in the hierarchical machine learning model that connects the input variables (i.e., chemical structures of drugs) and the output variables (i.e., PRRs for various ADRs), while allowing for a deeper mechanistic understanding of ADRs (**Fig. 1**).

### Prediction of Target, Enzyme, and Transporter Profiles from Chemical Fingerprints of Drugs

To predict the pharmacological profiles (i.e. TET profiles) from the chemical fingerprints (i.e. MACCS keys) of drugs, Random Forest Classifiers (RFCs) were constructed. The RFCs achieved high (>95%) accuracy across TET profile prediction (**Table 1**). The accuracies of these models were higher than other machine learning algorithms previously developed for the classification of drugs inhibiting a specific transporter [24]. For prediction of the entire TET profile, a testing accuracy of the RFC models is estimated to be 94.63% (=99.54% × 96.82% × 98.19 %; **Table 1**).

**Table 1.** Average training and testing accuracies for target, enzyme, and transporter prediction by the RFC models.

|  | Training Accuracy | Testing Accuracy |
| --- | --- | --- |
| Target | 99.69% | 99.54% |
| Enzyme | 98.74% | 96.82% |
| Transporter | 99.39% | 98.19% |

The TET prediction method presented in this study may address limitations of costly and often time-inefficient preclinical *in vitro* experiments to determine TET profiles [32]. Moreover, *in vitro* methods to assess drug’s action on transporters are not well established, presenting another limitation [33]. Other computational approaches, such as molecular docking, require the 3D chemical structures of TETs [34], which are often lacking in newly identified drug targets and many transporters [24]. Requiring no such 3D structural information, the RFCs presented here allow for accurate, thorough, and inexpensive evaluations of TET profiles of drugs — even those under the development stage.

### ADR prediction from Target, Enzyme, and Transporter Profiles of Drug Pairs

To predict ADRs of a drug pair from its TET profiles, Random Forest Classifier (RFC), Logistic Regression (LR), and Support Vector Machine (SVM) models were developed and evaluated using a 10-fold cross-validation. Compared to the RFC and LR models performing at mean classification accuracies (i.e. a fraction of a correctly classified ADR from a drug pair’s TET matrix) of 91.96% (**Fig. 2A**) and 86.63% (**Fig. 2B**), respectively, the SVM model outperformed with a greater mean classification accuracy of 95.73% (**Fig. 2C**).

**Fig. 2.**
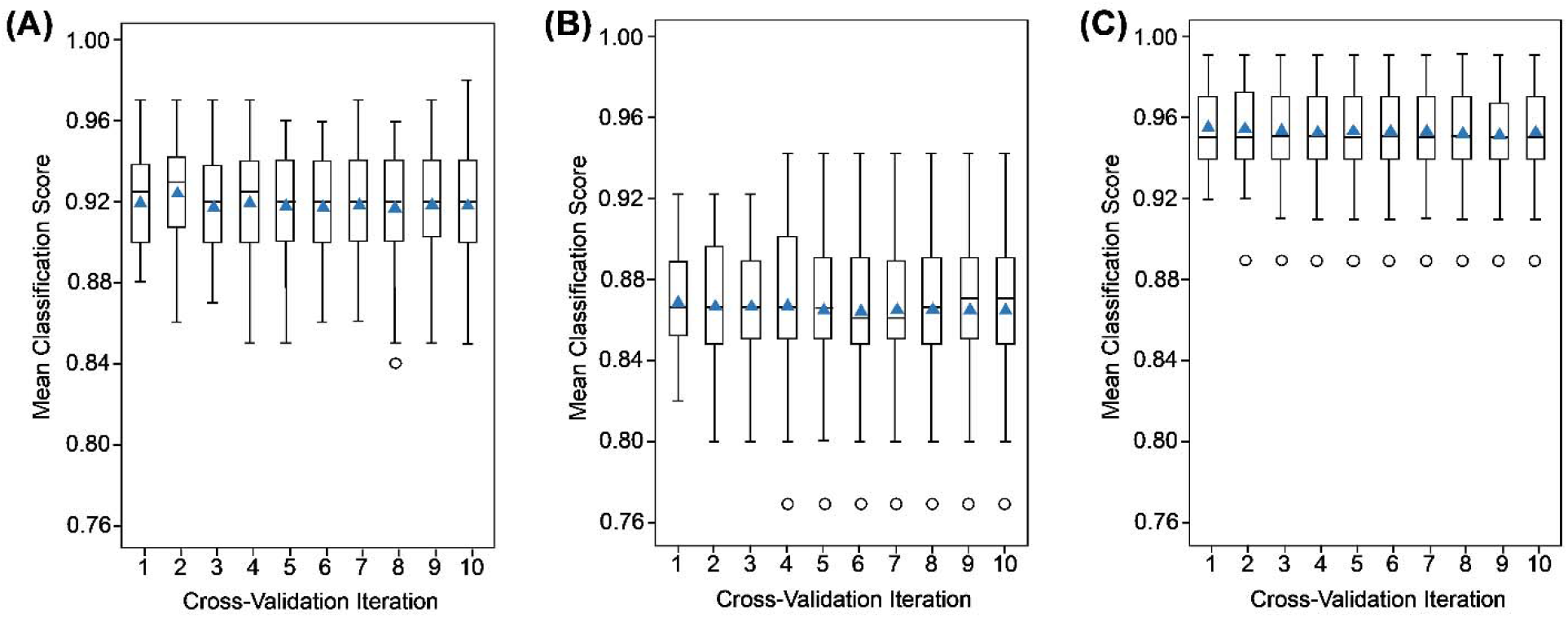
Repeated 10-fold cross-validation for (A) Random Forest Classifier (RFC), (B) Logistic Regression (LR), and (C) Support Vector Machine (SVM) models.

### Application of the SVM model for DDI-associated ADRs Involving Three Major Drugs

The SVM model was further tested for its robustness with DDIs involving three commonly prescribed drugs: levothyroxine, a synthetic hormone to treat hypothyroidism [35], omeprazole, a proton pump inhibitor for gastric acid-related disorders [36], and atorvastatin, an inhibitor of 3-hydroxy-3-methyl-glutaryl-CoA (HMG-CoA) reductase used for lowering lipids concentrations to treat hypercholesterolemia [37].

#### (1) Case study 1: Levothyroxine

To apply the SVM model for DDIs associated with levothyroxine, eptifibatide was chosen as the concomitant drug, since the co-administration of levothyroxine and a blood thinner (e.g., eptifibatide) was previously found to cause a bleeding-related ADR [38, 39] via inhibition of platelet aggregation [38, 40]. Levothyroxine has four major targets (integrin subunit αV (ITGAV), integrin subunit βIII (ITGB3), thyroid hormone receptor α (THRA), and thyroid hormone receptor β (THRB)), two metabolizing enzymes (cytochrome P450 (CYP) 2C8 (CYP2C8) and UDP-glucuronosyltransferase 1A1 (UGT1A1)), and nine transporters (ATP-binding cassette sub-family B member 1 (ABCB1), solute carrier (SLC) family 7 member 5 (SLC7A5), SLC16A2, solute carrier organic anion transporter (SLCO) 1A2 (SLCO1A2), SLCO1B1, SLCO1B3, SLCO1C1, SLCO2B1, and SLCO4A1; **Fig. 3A**). THRA and THRB are nuclear receptors for levothyroxine, regulating transcription of hormone-responsive genes (referred to as genomic actions) [41, 42]. A heterodimeric complex, ITGAV-ITGB3, consisting of ITGAV and ITGB3 is another receptor for levothyroxine [41, 42], mediating the drug’s nongenomic actions, such as the proliferation of endothelial cells [41, 42]. Eptifibatide’s TET profile contains only one target, ITGB3 (**Fig. 3A**), which is complexed with integrin subunit αIIb [43] to form a heterodimeric complex, ITGB3-ITGA2B, mediating platelet aggregation [40, 43]. The co-administration of eptifibatide, which inhibits ITGB3-ITGA2B’s binding to fibrinogen, can reduce platelet aggregation [38, 40]. Thus, the sharing of ITGB3 by levothyroxine and eptifibatide (**Fig. 3A**) may be responsible for ADRs, as is the case with other combinations of drugs binding to the same pharmacological targets [44].

**Fig. 3.**
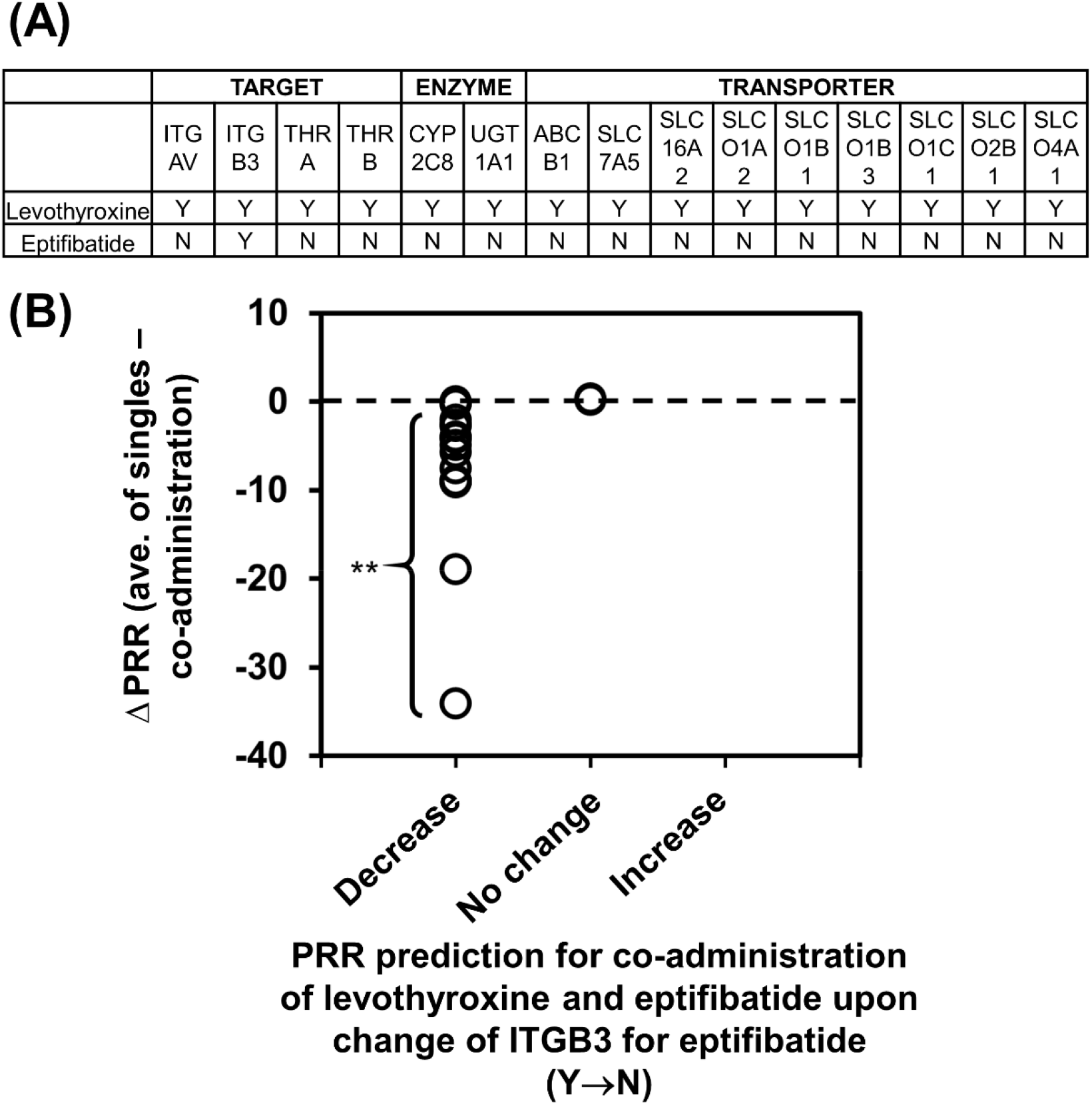
Adverse drug reactions associated with drug-drug interactions (DDIs) between levothyroxine and eptifibatide. **(A) Target, enzyme, and transporter (TET) profiles of levothyroxine and eptifibatide.** Y: the presence of a drug’s action on TETs. N: the absence of a drug’s action on TETs. **(B) Comparisons between** Δ**PRRs (as DDI indices) and PRR prediction upon removal of ITGB3 from eptifibatide’s TET profile for ADRs associated with the co-administration of levothyroxine and eptifibatide.** For a given ADR, the average PRR of single administrations of levothyroxine and eptifibatide – the PRR of their co-administration was calculated, and the PRR change for co-administration of levothyroxine and eptifibatide was predicted upon alteration of TET profiles of eptifibatide for integrin β-3 (ITGB3) from Y to N. Y: Inclusion of ITGB3 in eptifibatide’s TET profile. N: Removal of ITGB3 from eptifibatide’s TET profile. **: Outside of the 99% confidence interval of the “No change” group.

The predictive power of the SVM model, particularly in the role of the shared target (i.e. ITGB3) in DDI-associated ADRs, was evaluated through comparisons with statistical results. Briefly, the PRRs for various ADRs associated with the co-administration of levothyroxine and eptifibatide were calculated from statistical analyses of TWOSIDES v0.1. The PRRs for levothyroxine alone and eptifibatide alone were calculated similarly using OFFSIDES v0.1 [3]. Then, for a given ADR, its PRR for the co-administration of levothyroxine and eptifibatide was subtracted from the average PRR for the single administrations of levothyroxine and eptifibatide (i.e. Δ PRR = average PRR for single administrations of levothyroxine and eptifibatide – PRR for their co-administration). Highly negative values of this difference (e.g., Δ PRR < the 99% confidence interval of Δ PRRs for the “No Change” group) are indicative of strong DDIs. The calculated PRR differences were then compared with prediction results, which were obtained using the SVM model upon removal of the ITGB3 as a target from the TET profile of eptifibatide. The comparison result suggests that the risks of most ADRs associated with strong DDIs between levothyroxine and eptifibatide are predicted to decrease if the TET profile of eptifibatide lacks ITGB3 as a target, suggesting the critical role of shared ITGB3 in the DDIs (**Fig. 3B**).

#### (2) Case study 2: Omeprazole

For a subsequent analysis, clopidogrel, an antiplatelet drug for the treatment of cardiovascular diseases [45], was chosen as the concomitant drug with omeprazole. Omeprazole has two major targets (aryl hydrocarbon receptor (AHR) and potassium-transporting ATPase α chain 1 (ATP4A)), nine metabolizing enzymes (CYP1A1, CYP1A2, CYP1B1, CYP2C8, CYP2C9, CYP2C18, CYP2C19, CYP2D6, and CYP3A4), and three transporters (ABCB1, ABC subfamily C member 3 (ABCC3), and ABC subfamily G member 2 (ABCG2); **Fig. 4A**)

**Fig. 4.**
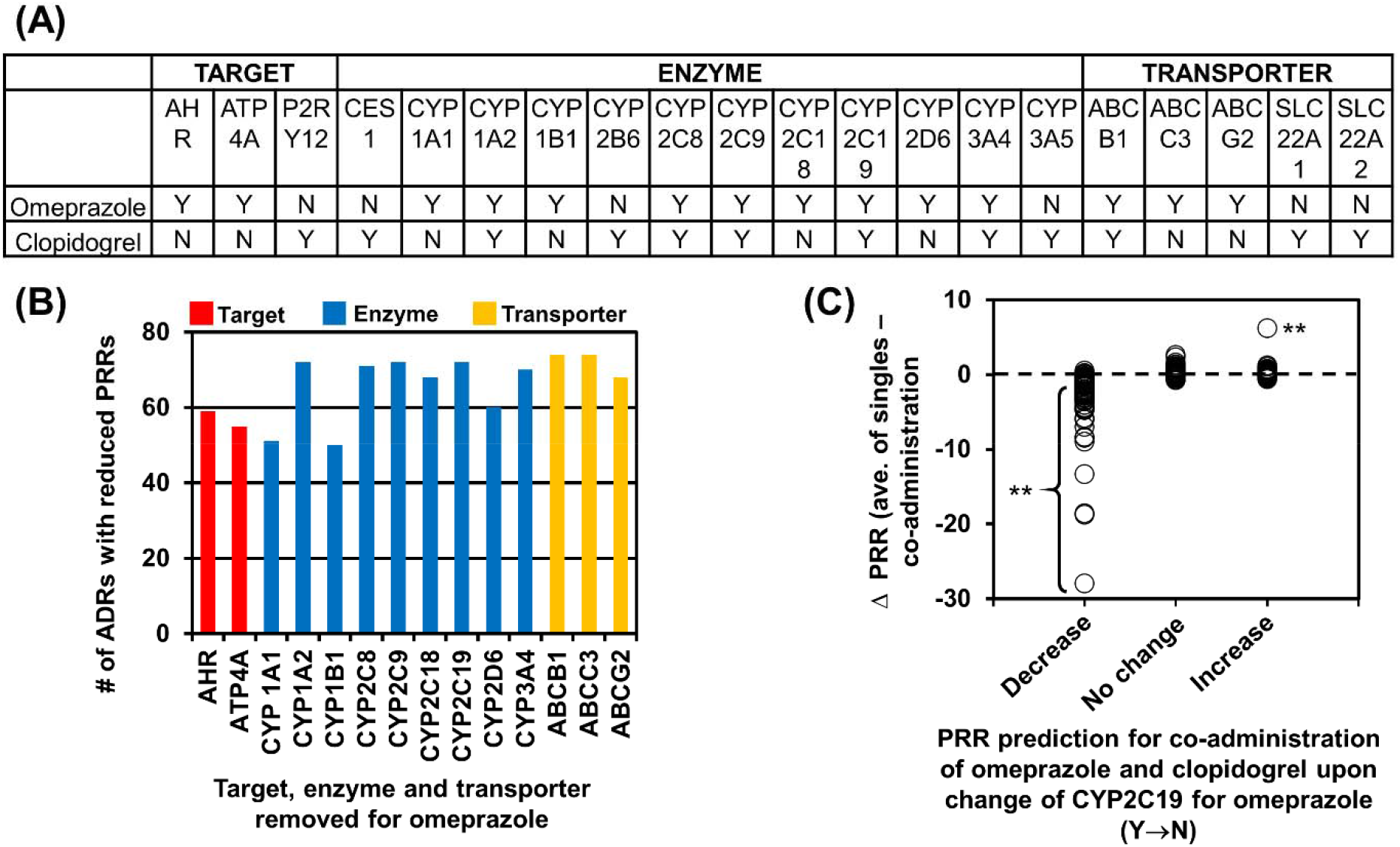
Adverse drug reactions associated with drug-drug interactions (DDIs) between omeprazole and clopidogrel. **(A) Target, enzyme, and transporter (TET) profiles of omeprazole and clopidogrel.**Y: the presence of a drug’s action on TETs. N: the absence of a drug’s action on TETs. **(B) The impacts of omeprazole’s TET profile on its DDI-associated ADRs with clopidogrel.**The PRR changes for ADRs associated with co-administration of omeprazole and clopidogrel were calculated using the SVM model when each of omeprazole’s TETs was removed. **(C) Comparisons between**Δ**PRRs (as DDI indices) and PRR predictions upon removal of CYP2C19 from omeprazole’s TET profile for ADRs associated with the co-administration of omeprazole and clopidogrel.**For a given ADR, the average PRR of single administrations of omeprazole and clopidogrel – the PRR of their co-administration was calculated, and the PRR change for co-administration of omeprazole and clopidogrel was predicted using the SVM model upon alteration of the TET profile of omeprazole for CYP2C19 from Y to N. Y: Inclusion of CYP2C19 in omeprazole’s TET profile. N: Removal of CYP2C19 from omeprazole’s TET profile. **: Outside of the 99% confidence interval of the “No change” group.

While omeprazole and clopidogrel share no same pharmacological targets, they have multiple enzymes and a single transporter in common (**Fig. 4A**). The concomitant use of omeprazole was found to lower the platelet inhibitory effects of clopidogrel [46, 47], increasing a risk of reinfarction [48, 49] and major cardiovascular events [50], compared to those receiving clopidogrel alone. Accordingly, the FDA has recommended not using omeprazole together with clopidogrel unless absolutely required [47].

To identify the major pharmacological determinants responsible for DDIs between omeprazole and clopidogrel, each of the targets, enzymes and transporters was removed one at a time from the TET profile of omeprazole and the effects of each removal on the DDI-associated ADRs were predicted using the SVM model developed in this study. The result from this analysis showed that CYP1A2, CYP2C8, CYP2C9, CYP2C19, CYP3A4, ABCB1, and ABCC3 may play key roles in DDIs between omeprazole and clopidogrel (**Fig. 4B**).

Consistent with this result, CYP2C19 was previously identified as a key enzyme to mediate DDIs between omeprazole and clopidogrel [47]. For its anti-platelet aggregation effect, clopidogrel needs to be converted by CYP2C19 to an active metabolite [51, 52], which prevents activation of P2RY12 required for platelet activation and aggregation [53]. Thus, omeprazole, an inhibitor of CYP2C19 [47, 54], can prevent the biotransformation of clopidogrel required for efficacy, causing DDI-associated ADRs [46–49]. The analysis also suggests the possible involvement of other enzymes and transporters in DDIs between omeprazole and clopidogrel (**Fig. 4B**), as supported by previous reports. For example, the metabolic activation of clopidogrel was also found to be mediated by CYP3A4 [55, 56]. In addition, the high likelihood of CYP1A2 mediating DDIs involving omeprazole was previously proposed based on omeprazole’s ability to induce CYP1A2 activity [57, 58], though still under debate [47, 59, 60]. Omeprazole is a weak inhibitor of CYP2D6 relative to CYP2C19 and CYP3A4 [47], making CYP2D6-mediated DDIs less likely. An efflux transporter, ABCB1, may be an active player in these DDIs, as omeprazole interferes with the efflux of other drugs (e.g., digoxin [61] and nifedipine [62]) by ABCB1 [47].

Out of these enzymes and transporters, CYP2C19 was chosen as a key enzyme for subsequent comparative analyses. For this examination, Δ PRR (= an average PRR of omeprazole alone and clopidogrel alone – a PRR for the co-administration of omeprazole and clopidogrel) was calculated as a DDI index for each ADR, as described above. Δ PRR values were then compared with the PRR changes predicted by the SVM model when CYP2C19 was removed from the TET profile of omeprazole. Overall, the predictions and the calculations were in good agreement for ADRs significantly associated with DDIs (either negatively or positively, as judged by Δ PRR relative to the 99% interval of Δ PRRs for the “No change” group) between omeprazole and clopidogrel (**Fig. 4C**), supporting the role of CYP2C19 in their DDIs, as described elsewhere [47]. The comparative result also indicates that correct predictions of drug pairs with little to no DDIs (that is, Δ PRR ~ 0) are difficult with this model.

#### (3) Case study 3: Atorvastatin

To further validate the SVM model, a similar computational approach was applied to atorvastatin for its well-known DDI-associated ADR, myopathy [63–66]. Atorvastatin has five major targets (AHR, dipeptidyl peptidase 4 (DPP4), histone deacetylase 2 (HDAC2), 3-hydroxy-3-methylglutaryl-coenzyme A reductase (HMGCR), and nuclear receptor subfamily 1 group I member 3 (NR1I3)), ten metabolizing enzymes (CYP2B6, CYP2C8, CYP2C9, CYP2C19, CYP2D6, CYP3A4, CYP3A5, CYP3A7, UGT1A1, and UDP-glucuronosyltransferase 1A3 (UGT1A3)), and ten transporters (ABCB1, ATP-binding cassette sub-family B member 11 (ABCB11), ATP-binding cassette sub-family C member (ABCC) 1 (ABCC1), ABCC2, ABCC4, ABCC5, SLCO1A2, SLCO1B1, SLCO1B3, and SLCO2B1; **Fig. 5A**). Atorvastatin’s actions on multiple TETs indicate its pharmacological complexity.

**Fig. 5.**
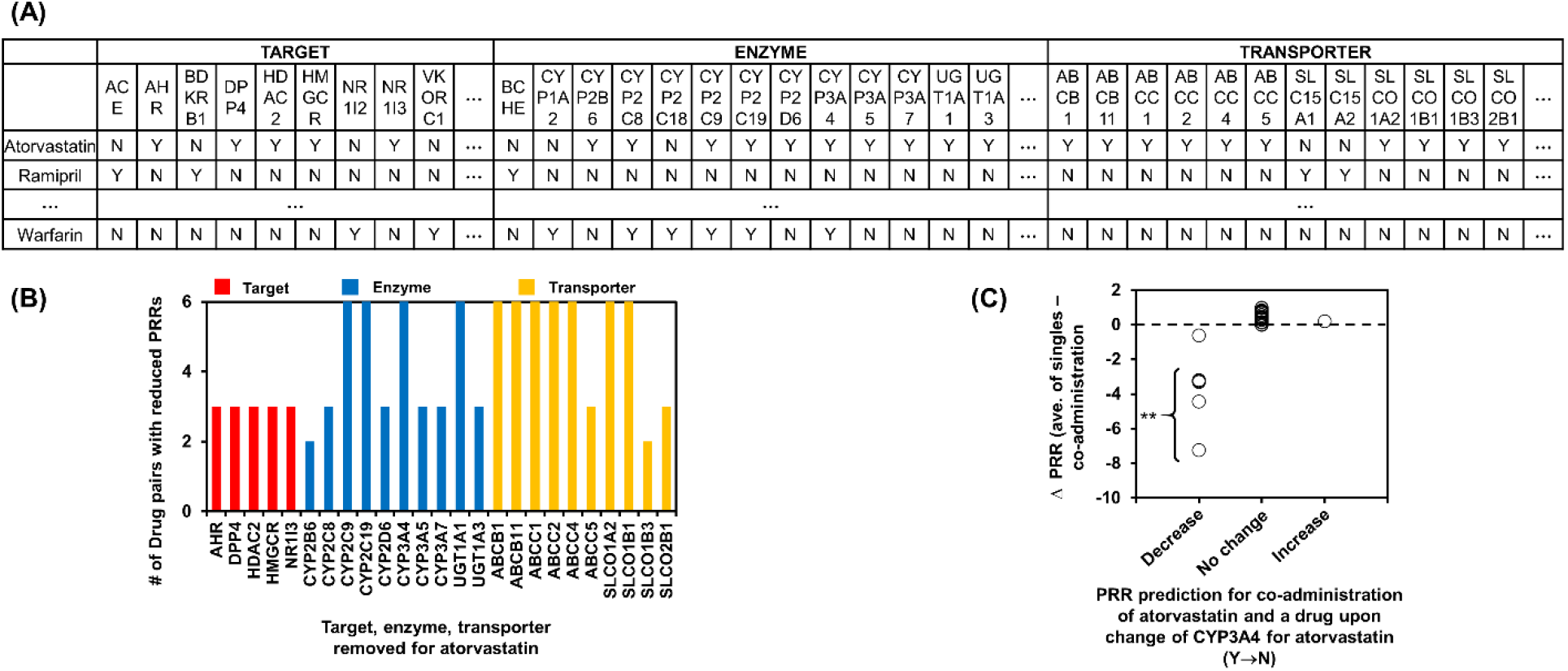
Myopathy associated with drug-drug interactions (DDIs) involving atorvastatin. **(A) Target, enzyme, and transporter (TET) profiles of atorvastatin and concomitant drugs, such as ramipril and warfarin.** Y: the presence of a drug’s action on TETs. N: the absence of a drug’s action on TETs. **(B) The impacts of atorvastatin’s TET profile on its DDI-associated ADR of myopathy with various concomitant drugs.**The PRR changes for myopathy associated with co-administration of atorvastatin and other drugs were calculated using the SVM model when each of atorvastatin’s TETs was removed. **(C) Comparisons between** Δ**PRRs (as DDI indices) and PRR predictions upon removal of CYP3A4 from atorvastatin’s TET profile for myopathy associated with the co-administration of atorvastatin and a concomitant drug.**For myopathy, the average PRRs of single administrations of atorvastatin and a concomitant drug – the PRRs of their co-administration were calculated, and PRR changes for the co-administration of atorvastatin and the drug were predicted using the SVM model upon alteration of the TET profile of atorvastatin for cytochrome P450 3A4 (CYP3A4) from Y to N. Y: Inclusion of CYP3A4 in atorvastatin’s TET profile. N: Removal of CYP3A4 from atorvastatin’s TET profile. **: Outside of the 99% confidence interval of the “No change” group.

The SVM model was used to derive TETs important in atorvastatin-induced myopathy through predicted PRR changes of drug pairs upon removal of each of the targets, enzymes, and transporters from atorvastatin’s TET profile. The SVM model predicted the importance of CYP2C9, CYP2C19, CYP3A4, UGT1A1, ABCB1, ABCB11, ABCC1, ABCC2, ABCC4, SLCO1A2, and SLCO1B1 in atorvastatin DDI-associated myopathy (**Fig. 5B**). Consistent with this result, the co-administration of drugs that are either inhibitors or substrates of CYP3A4 was found to decrease the metabolism of atorvastatin [67]. As a result, the plasma concentration of atorvastatin increases, leading to the onset of ADRs [67], including myopathy [66]. In addition, polymorphisms in the *CYP2C19*, *UGT1A1*, *ABCB1* and *SLCO1B1* genes are associated with systemic exposure of atorvastatin, an important risk factor for myopathy [68, 69]. Drugs inhibiting SLCO1B1 and ABCB1, most of which are CYP3A4 inhibitors [70], can cause DDIs with atorvastatin [70]. While the crucial role of CYP2C8 in DDIs involving other statins (e.g., simvastatin and lovastatin) has been documented [70, 71], the involvement of this enzyme in atorvastatin-mediated DDIs remains unclear.

The model was then tested through comparisons between the predicted and calculated PRR changes of drug combinations for myopathy. Out of the identified key molecules, CYP3A4 was chosen for further analyses due to its direct involvement in the onset of myopathy associated with DDIs involving atorvastatin, as reported previously [72, 73]. PRRs of drugs with higher degrees of DDIs (as judged by Δ PRR relative to the 99% confidence interval of Δ PRRs for the “No Change” group) with atorvastatin were predicted to decrease upon removal of atorvastatin’s CYP3A4 interaction (**Fig. 5C**), consistent with the literature reports on the importance of this CYP enzyme in atorvastatin-mediated myopathy [72, 73]. Similar to the results with omeprazole (case study 2), the accurate prediction of PRR changes for drug combinations with Δ PRR~ 0 was difficult (**Fig. 5C**).

Overall, the results obtained with levothyroxine, omeprazole and atorvastatin in this study demonstrate the high applicability of the machine learning model for predicting DDI-associated ADRs and providing underlying pharmacological insight.

### Model Application for Interstitial Lung Disease Involving DDIs with Atorvastatin

Motivated by its high prediction power, the model developed in this study was applied to a rare yet life-threatening ADR, interstitial lung disease (ILD), associated with DDIs involving atorvastatin. The PRR of the single administration of atorvastatin for ILD was calculated from OFFSIDES v0.1 to analyze its statistical associations to ILD. A similar statistical analysis was extended to drug pairs containing atorvastatin for ILD. Δ PRRs (i.e. DDI indices) were calculated and plotted with PRRs of the co-administration for ILD (**Fig. 6**).

**Fig. 6.**
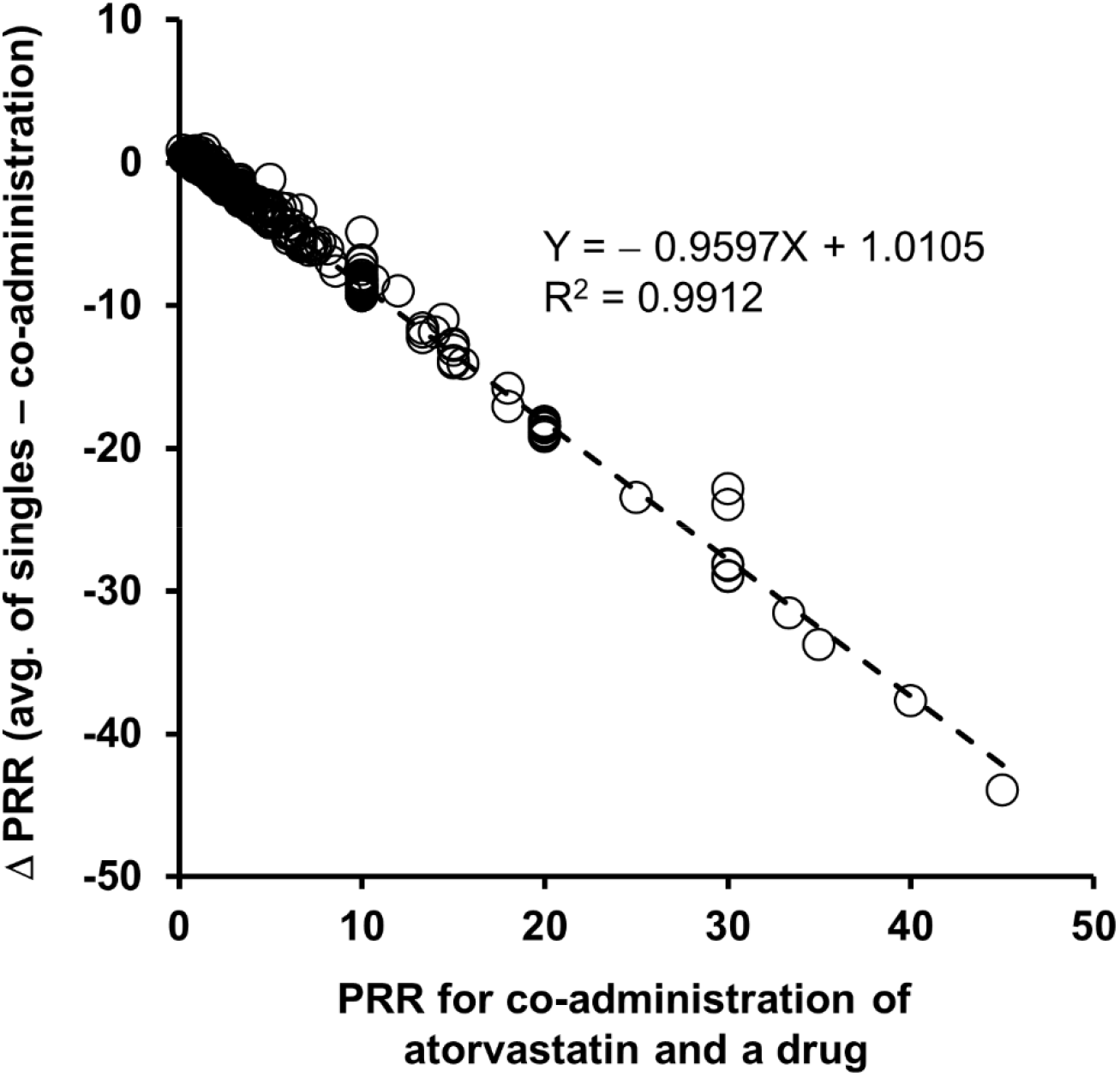
Drug-drug interactions between atorvastatin and concomitant drugs for interstitial lung disease (ILD). For ILD, Δ PRR (= the average PRR of single administrations of atorvastatin and the concomitant drug – the PRR of their co-administration) was calculated, and plotted with the PRR for their co-administration.

The PRR of atorvastatin alone for ILD was 1.01, a value indicative of a statistically neutral association between atorvastatin and ILD. Between ΔPRRs and PRRs for the co-administration of atorvastatin, a strong negative linear relationship was detected (**Fig. 6**). The implication is that drug pairs of atorvastatin and concomitant drugs reported with high risks of ILD are due to DDIs between the drugs. When calculated from the linear regression line, Δ PRR is 0.0508 (~ 0, no DDI) for drug pairs showing no associations with ILD (i.e., PRR = 1; **Fig. 6**), as expected, validating this analysis.

To identify important TETs in ILD associated with DDIs involving atorvastatin, the PRR changes of drug pairs were predicted by the model upon removal of each of the targets, enzymes, and transporters from atorvastatin’s TET profile (**Fig. 7A**). Among atorvastatin’s five targets, AHR, DPP4, HDAC2, HMGCR and NR1I3, the importance of DPP4 was minimal (**Fig. 7A**). In this analysis, different metabolizing enzymes seemed equally important, suggesting that DDI-associated ILD involving atorvastatin may be mediated by a set of multiple enzymes, which may be responsible for previous contradictory findings on the role of metabolizing enzymes in this type of ADR [74, 75]. Among transporters, ABCB11, SLCO1B1 and SLCO1B3, all of which are primarily expressed in liver [30], were found to be important. ABCB11, a primary transporter of bile salts [76], was found to be involved in the biliary excretion of statins [77]. SLCO1B1 and SLCO1B3 are responsible for the uptake of atorvastatin into hepatocytes [78, 79]. Thus, the removal of these three transporters from atorvastatin’s TET profile may increase its plasma concentration, increasing risks of ILD [78, 79].

**Fig. 7.**
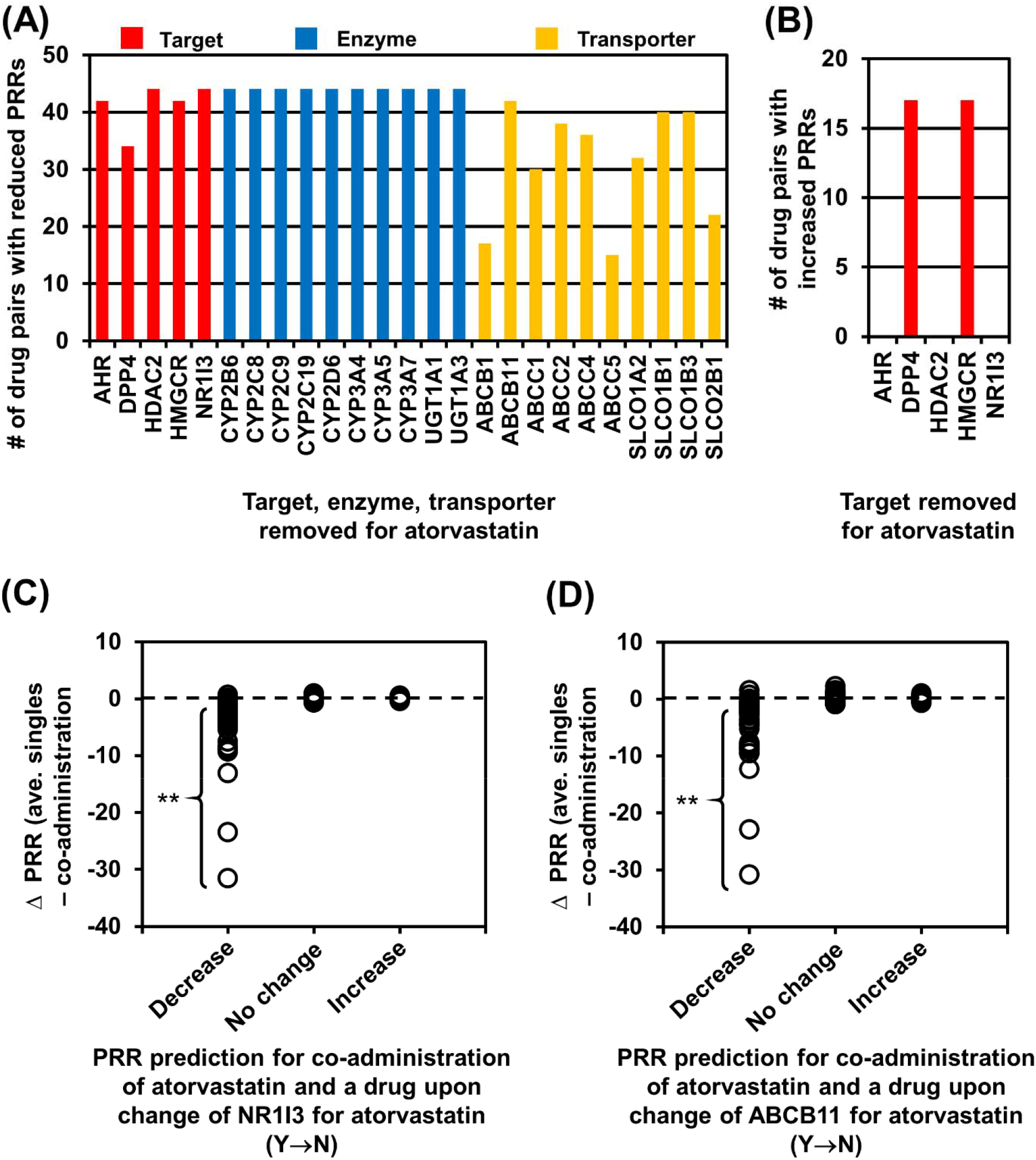
Interstitial lung disease (ILD) associated with drug-drug interactions (DDIs) involving atorvastatin. **(A) The impacts of atorvastatin’s TET profile on its DDI-associated ILD with various concomitant drugs.**The PRR changes for ILD associated with the co-administration of atorvastatin and other drugs were calculated using the SVM model when each of atorvastatin’s TETs was removed. **(B) The number of drug pairs containing atorvastatin with a predicted increase in PRRs for ILDs when each of atorvastatin’s targets was removed. (C) Comparisons between** Δ**PRRs (as DDI indices) and PRR predictions upon removal of (A) NR1I3 and (B) ABCB11 from atorvastatin’s TET profile for ILD associated with the co-administration of atorvastatin and a concomitant drug.**For ILD, the average PRR of single administrations of atorvastatin and a concomitant drug– the PRR of their co-administration was calculated, and the PRR changes for the co-administration of atorvastatin and the drug were predicted using the SVM model upon alteration of the TET profile of atorvastatin for (A) NR1I3 and (B) ABCB11 from Y to N. Y: Inclusion of (A) NR1I3 and (B) ABCB11 in atorvastatin’s TET profile. N: Removal of (A) NR1I3 and (B) ABCB11 from atorvastatin’s TET profile. **: Outside of the 99% confidence interval of the “No change” group.

To further distinguish among the five targets, a similar procedure was conducted to calculate the number of drug pairs with predicted increases in PRRs for ILD when a target was removed from atorvastatin’s TET profile (**Fig. 7B**). Interestingly, when atorvastatin’s action on HMGCR and DPP4 became nullified, PRRs for ILD further increased (**Fig. 7B**). The implication is that when its binding to HMGCR and DPP4 becomes ineffective, atorvastatin may bind to the other three targets more strongly, increasing risks of DDIs. No such PRR increases were observed with the removal of the other three targets (**Fig. 7B**). Thus, AHR, HDAC2, and NR1I3 were identified as important targets for DDI-associated ILD involving atorvastatin.

To validate these computational results, two key molecules, NR1I3 and ABCB11, which were identified by the SVM model in DDI-associated ILD with atorvastatin, were used for further analyses. For this examination, Δ PRRs (the average PRR for single administrations of atorvastatin and a concomitant drug – the PRR for their co-administration) were calculated and compared with PRR predictions by the SVM model for the drug pairs upon the removal of NR1I3 (**Fig. 7C**) and ABCB11 (**Fig. 7D**) from the TET profile of atorvastatin. In these analyses, PRRs for ILD were predicted to decrease with most drug pairs involving significant DDIs, when judged by Δ PRR < the 99% confidence interval of Δ PRRs for the “No change” group (**Fig. 7C-D**), supporting the critical roles of NR1I3 and ABCB11 in DDI-associated ILD involving atorvastatin.

A few potential pathological pathways underlying DDI-associated ILD involving a high plasma concentration of atorvastatin were determined around the three important targets — AHR, NR1I3, and HDAC2 — identified in this study. In this analysis, only genes/proteins significantly expressed in lung, as recorded in the NCBI Gene database were [30] considered. The analyses revealed the high likelihood that atorvastatin binding to AHR, NR1I3, and HDAC2 may cause ILD through a major ILD mechanism — the dysregulation of surfactant production and homeostasis [80, 81]. Many interactors, such as SP1 transcription factor and estrogen receptor 1 (ESR1), are highly interconnected in the pathways around AHR, NR1I3, and HDAC2 (**Fig. 8A-B**). Genes/proteins important in surfactant metabolism also create a strong network (**Fig. 8A-B**). Thus, any impact from AHR, NR1I3 and HDAC2 can be amplified, influencing one another in these pathways.

**Fig. 8.**
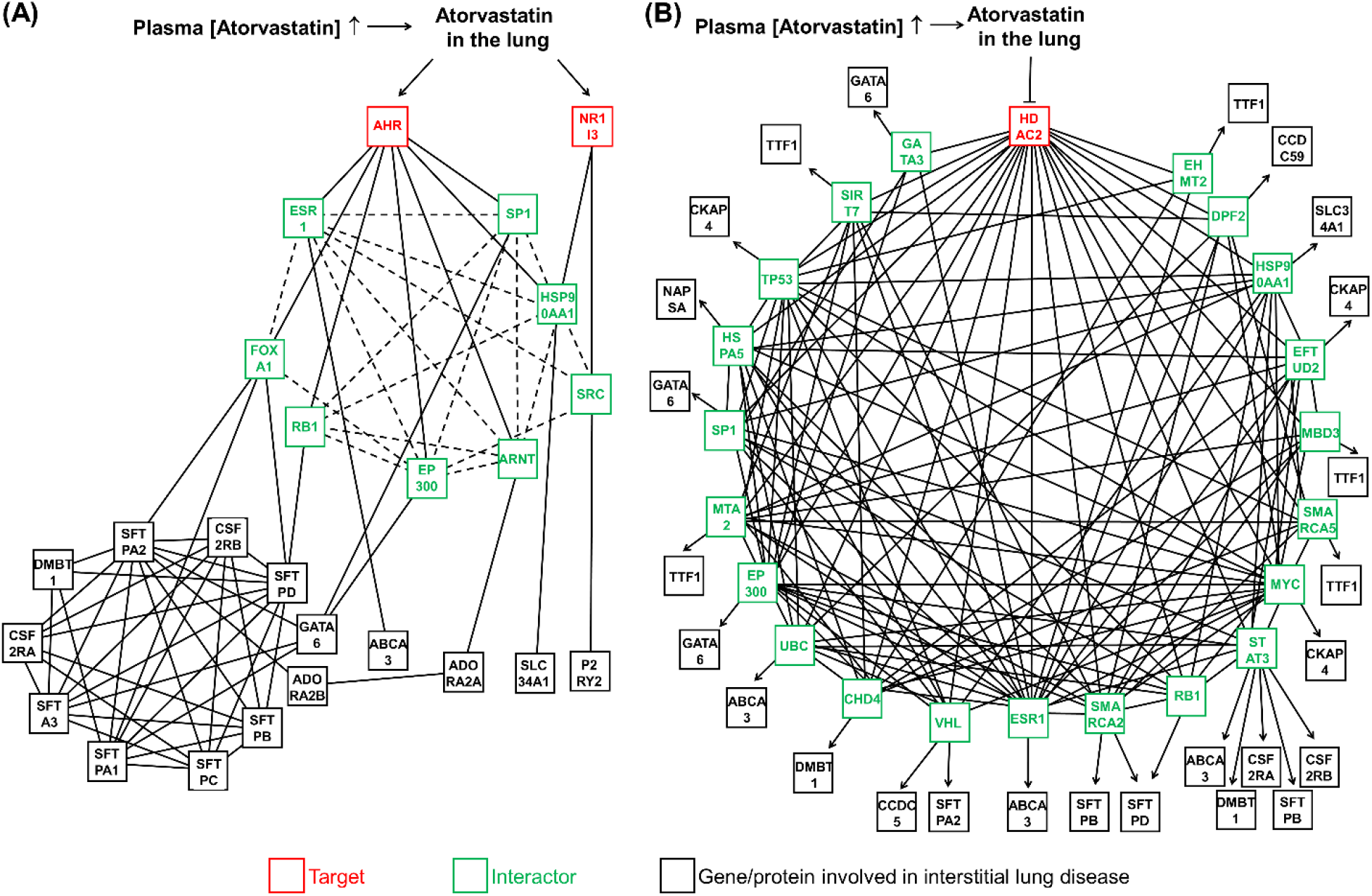
Pathway analyses for the enhanced risk of ILD, associated with DDIs involving atorvastatin created around. **(A) AHR and NR1I3 and (B) HDAC2.** Interactions among genes/proteins were determined using an array of bioinformatics databases, including BioGRID, Proteomics DB, STRING, CORUM and Reactome.

Literature survey identified a few plausible routes atorvastatin can take to cause ILD. AHR is a transcription factor inducible by aromatic hydrocarbon-based xenobiotics, such as atorvastatin [82, 83]. Upon binding to a ligand, AHR is complexed with aryl hydrocarbon receptor nuclear translocator (ARNT; **Fig. 8A**) [82, 83]. The AHR/ARNT complex can then induce expression of AHR’s target genes, which code for enzymes and transporters required for xenobiotic metabolism [82, 84]. Activated AHR inhibits estrogen receptor (ESR1) activity [83], by redirecting ESR1 away from ESR1 target genes [85], such as ATP-binding cassette sub-family A member 3 (*ABCA3*; **Fig. 8A**) [86]. ABCA3 plays a critical role in the formation of pulmonary surfactant by transporting phospholipids from the endoplasmic reticulum to a surfactant storage organelle in type II epithelial cells [87, 88]. Thus, the binding of atorvastatin to AHR may cause pulmonary surfactant metabolism dysfunction by downregulating the *ABCA3* gene via inhibition of ESR1 activity [89]. Different interactors (e.g., histone acetyltransferase p300 (EP300)) may be involved in ILD, amplifying the effects of AHR through networks of other interactors and ILD genes/proteins.

In addition, NR1I3 is a nuclear receptor that mediates transcriptional activation of target genes required for the metabolism and elimination of xenobiotics [82, 90, 91], such as CYP2B6 and CYP3A4 [90, 92]. Upon the binding of xenobiotics, NR1I3 is dephosphorylated for nuclear translocation and transactivation [93], which requires reduced SRC kinase activity [94]. On the other hand, P2Y purinoceptor 2 (P2RY2), a G protein-coupled receptor, activates SRC [95, 96] in order to promote surfactant secretion from alveolar type II cells [97]. Thus, the binding of atorvastatin to NR1I3 [98] can dysregulate normal surfactant secretion via interference with SRC kinase activity.

Atorvastatin inhibits HDAC2 [99]. The connections between HDAC2 and ILD genes/proteins are highly interconnected, also sharing many interactors with AHR and NR1I3, suggesting the existence of many different paths that cause ILD from HDAC2 inhibition (**Fig. 8B**). Interestingly, the binding of atorvastatin to HDAC2 may be related to atorvastatin’s beneficial effect against cancer [100]. HDAC2 plays a key role in the epigenetic regulation of gene expression in cancer [101] and HDAC2 inhibitors (e.g., atorvastatin [99]) can display anti-cancer activities [101]. The anticancer effect of atorvastatin was also statistically analyzed. In this analysis, combinations of atorvastatin and a drug that have PRRs > 1 for ILD were identified and their PRRs for lung cancer were also calculated. None of combinations of atorvastatin and concomitant drugs that had PRRs > 1 for ILD had PRRs > 1 for lung cancer, supporting atorvastatin’s anti-cancer effects. Overall, the model-based computational examinations and pathway analyses revealed key molecules important in DDI-associated ILD involving atorvastatin and proposed underlying pathological pathways.

## CONCLUSIONS

This study presented a novel computational approach to accurately predict the occurrences of ADRs using a machine learning model consisting of hierarchically structured classifiers. The hierarchical model presented here addresses the limitations of conventional models relying on drug similarity for the prediction of ADRs. The method developed here is based on TET profile-dependencies of ADRs derived from drugs’ chemical structures, requiring no high chemical similarity of drugs. Given basic structural characteristics of drugs, this hierarchical model integrating the RFCs for TET profile prediction and the SVM for specific DDI-associated ADRs can accurately predict ADRs with an overall ~91% (=94.63% for TET prediction × 95.73% for ADR prediction) accuracy. As DDIs typically appearing as various forms of ADRs have been another primary issue in past predictions of DDI-associated ADRs [102], the presented model deconvolutes this complexity, as judged by its accurate prediction of various ADRs for any drug pair. In addition, pharmacological insight offered by the hierarchical model was successfully connected to pathway analyses underlying ADRs, making the described computational approach powerful for not only predicting the occurrence of DDI-associated ADRs but also enhancing mechanistic understandings.

Notably, the constructed model can accurately predict TET profiles and DDI-associated ADRs from most basic information of drugs - chemical structures – for any pair. Thus, the model presented in this study can also be used for drug design. For example, the MACCS keys descriptors can be manipulated and inputted into the hierarchical model to identify a drug’s key structural characteristics that increase a risk of ADRs. Once the structural hot spots are identified, an array of drug variants with different chemical moieties at the locations can be designed and evaluated for DDIs and ADRs prior to synthesis. As a result, many drugs can readily be evaluated for their potential DDIs in advance, avoiding costly preclinical and clinical tests. Thus, the hierarchical model developed is anticipated to pave new way to enhance drug safety and reduce drug development costs.

## Notes

### Competing Interest Statement

The authors have declared no competing interest.

